# Hybrid Synthesis of bioplastics polyhydroxybutyrate from carbon dioxide

**DOI:** 10.1101/2022.09.30.510340

**Authors:** Jie Zhang, Dingyu Liu, Yuwan Liu, Huanyu Chu, Jian Cheng, Haodong Zhao, Shaoping Fu, Liu Huihong, YuE Fu, Yanhe Ma, Huifneg Jiang

**Author notes:** These authors contributed equally to this work (Jie Zhang, Dingyu Liu). Corresponding author. Email: Huifeng Jiang,; Yanhe Ma,.

## Abstract

The accelerating environmental crisis has intensified the demand for switching from traditional economy to a renewable one with a reduced carbon footprint. Here we reported a hybrid system, coupling chemical process of CO_2_ hydrogen reduction and biological process for polyhydroxybutyrate (PHB) synthesis, that utilized CO_2_ as a raw material to produce PHB in vitro. The synthetic pathway of PHB was optimized by screening more efficient methanol oxidases, high activity mutants of glycolaldehyde synthase and coordinating enzyme dosages in the pathway, which achieved the carbon yield of 93.6% for producing PHB from methanol. Finally, by combining with the chemical process from CO_2_ to methanol, a scaling-up bio-system was performed to convert CO_2_ into PHB, yielding 5.8 g/L with the productivity of 1.06 g^-1^L^-1^h^-1^. This approach represents a promising carbon-neutral way to produce biodegradable plastics.

## Introduction

The atmospheric CO_2_ concentration are increasing at an alarming rate, resulting in dire climate change effects. The use of CO_2_ as feedstock delivers environmentally sustainable, carbon-negative manufacturing of chemicals and materials.[1,2]. Polyhydroxybutyrate (PHB) is a class of biodegradable plastics, a well-recognized renewable and petroleum-based plastics substitute. Over the last 30 years, advances in metabolic engineering have enabled construction of microbial cell factories and cell-free systems for PHB production from sugar-based feedstock[3]. Although great success has been achieved, large-scale production and commercial applications remain limited due to manufacturing expense, approximately 50% of which are generated from raw materials[4]. Therefore, the utilization of CO_2_ as a source material in the synthesis of biodegradable plastics PHB offers a win-win strategy to both decrease the CO_2_ emissions and solve plastic pollution.

Photosynthetic carbon fixation is the main natural CO_2_ fixation process[5]. However, the titers and productivities of PHB in the engineered cyanobacteria were relatively low to unable economic feasibility and suffers from very low solar energy efficiency and slow growth rate[6, 7]. The *Cupriavidus necator* is also a very attractive autotroph platform with half a century of research history, and it can produce PHB from CO_2_, O_2_ and H_2_ at a rate of up to 1.55 g/L/h[8]. However, the large-scale production of PHB by mixed-gas fermentation presents a serious explosive risk. Despite significant efforts in chemistry, biology, materials, and engineering to develop CO_2_ fixation and utilization, comprehensive solutions remain to be explored.

Here we designed a hybrid system, coupling photovoltaic hydrogen production, CO_2_ hydrogenation with chemoenzymatic PHB synthesis, that drove an artificial environmentally friendly carbon-negative process for carbon fixation into PHB. A space-coupled system with excellent carbon fixation rate and molar conversion efficiency were obtained. In a chemoenzymatic process, carbon molar yield of 93.6% was achieved by a two-step pathway for producing PHB from methanol. Finally, a scaling-up hybrid system was performed to convert CO_2_ into PHB, yielding 5.8 g/L with the productivity of 1.06 g^-1^L^-1^h^-1^.

## Results

### Construction of hybrid system for PHB synthesis from CO_2_

Combining the captured CO_2_ with hydrogen produced from renewable energy to synthesize green methanol[9], has great potential to provide precursors for one carbon bio-manufacturing. The Synthetic Acetyl-CoA (SACA) pathway, a carbon-conserved and ATP-independent acetyl-CoA synthesis pathway, was applied to convert one carbon to acetyl-CoA[10]. In fact, the carbon yield of PHB production is determined by the yield of acetyl-CoA in the synthetic pathway, which usually was limited by inherent carbon loss during the natural CO_2_ assimilation pathway. Integrating with the synthetic pathway from acetyl-CoA to PHB, a powerful driving system for energy conversion and material metabolism is promised to produce PHB from CO_2_ with high efficient and carbon yield (Fig. 1). The technology of CO_2_ hydrogenation was employed to drive CO_2_ fixation to provide one-carbon (C1) unit[11]. Methanol was served as intermediate to bridge chemocatalysis and biosynthesis in this work. We coupled methanol oxidation reaction with the SACA pathway to convert C1 unit into acetyl-CoA[10], which is then converted into PHB by the synthesis pathway of PHB[12]. All reactions and the standard Gibbs free energy change (ΔG’) of each enzymatic reaction are listed in Table S2. This pathway efficiently converted CO_2_ to PHB with theoretical carbon yield of 100%, which displayed a significant atom-economic advantage compared to the natural metabolic pathway.

**Fig. 1.**
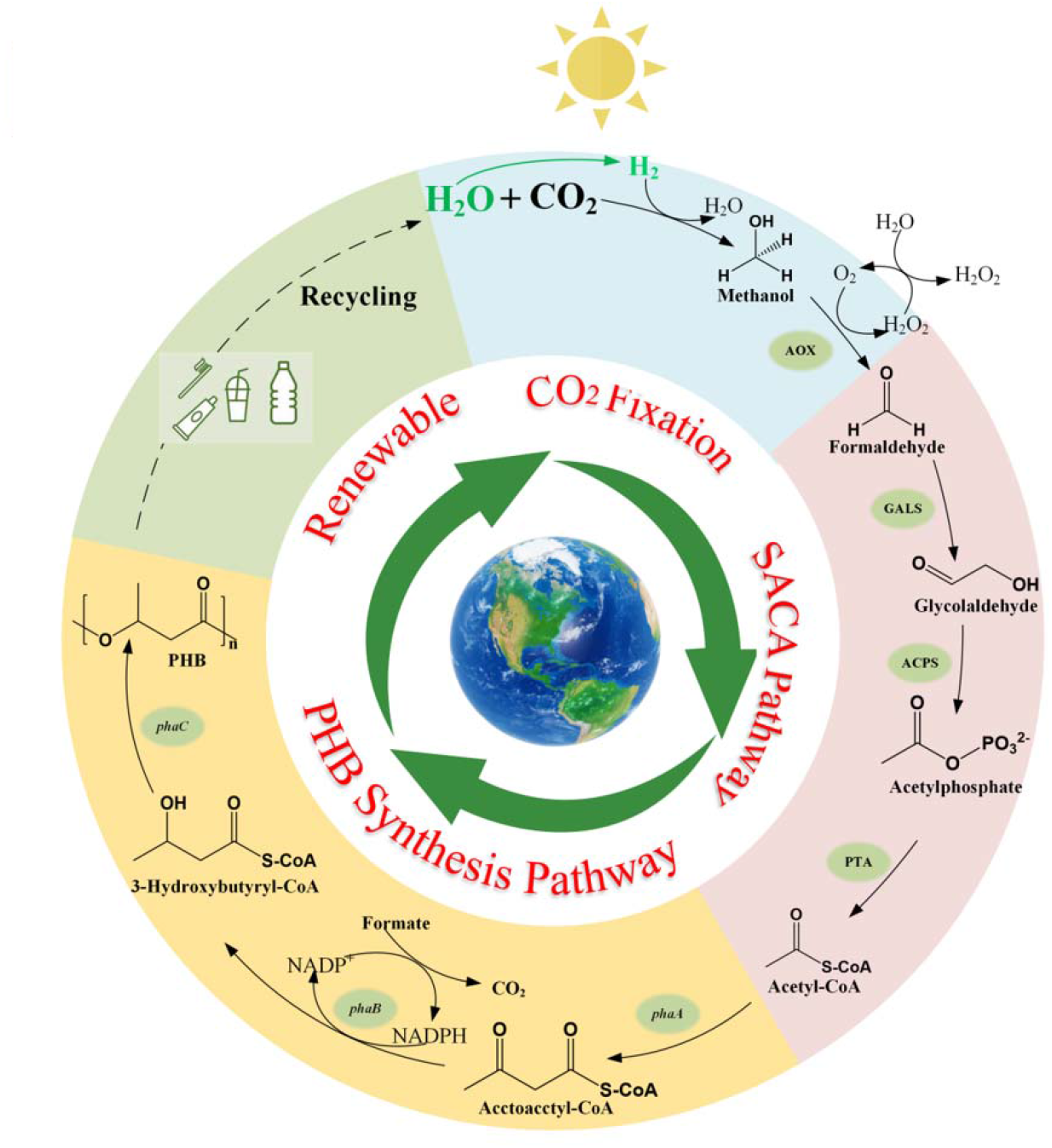
Design of hybrid system for producing PHB from CO_2_. Inner circle:A complete closed-loop carbon cycle for PHB synthesis to degradation. The SACA pathway indicates Synthetic Acetyl-CoA pathway. Green labelled H_2_O and H_2_ indicates the process of hydrogen production by water electrolysis. Outer circle: schematic of hybrid system, coupling photovoltaic hydrogen production, CO_2_ hydrogenation with chemoenzymatic PHB synthesis and environmental degradation. The individual modules were colored. Auxiliary enzymes and chemicals are indicated.

To prove this design, we used 20 mM methanol as substrate to synthesize PHB. However, one-pot enzymatic synthesis enabled inefficient production of PHB, only representing approximately 10% of the theoretical molar conversion efficiency of carbon (Supplementary Fig. 1). Acetate produced by the spontaneous hydrolysis of acetyl-phosphate was the main by-product in the pathway[13], since the carbon flux was kinetically trapped at acetyl-phosphate due to subsequent thermodynamically unfavourable reaction. At the same time, NADPH, the downstream driving force of thermodynamically unfavourable reaction, was also oxidized by catalase (CAT)[14] and methanol oxidase (AOX) (Supplementary Fig. 2). Thus, the acetyl-phosphate preferred to be convert into acetate. In addition, formaldehyde is toxic for many enzymes[15]. In order to run well the desired enzymatic cascade, we proposed to redivide the pathway into three modules, the module I containing all chemical reactions from CO_2_ to methanol; the module II including biological reactions from methanol to glycolaldehyde and the module III involving enzymic cascade from glycolaldehyde to PHB.

### Dynamic modulation of module II

Methanol was converted into glycolaldehyde via condensation of formaldehyde, an enzymatic cascade reaction containing AOX, CAT and glycolaldehyde synthase (GALS) (Fig.2a). In exception to the physiological substrate methanol, the AOX can also oxidize formaldehyde into formate due to the hydration of formaldehyde in aqueous solution[16]. Considering the principle of atomic economy, we proposed to employ formate to regenerate NADPH by formate dehydrogenase (FDH) for module III. In fact, if the ratio of formaldehyde/formate is 4:1, the generated NADPH is perfect for the synthesis of PHB. Thus, we screened AOX genes with significant formaldehyde specificity based on product distribution in the case of methanol as a substrate (Fig.2b). The PcAOX (from *Phanerochaete chrysosporium*) was finally selected for glycolaldehyde synthesis module, which was capable of balancing carbon fluxes of NADPH regeneration and PHB synthesis via formaldehyde[17]. The percentages of formaldehyde and formate were 84% and 16%, which almost achieved the theoretical stoichiometry for converting methanol to PHB (Fig. 2b). Finally, the glycolaldehyde synthesis module enabled 8 mM glycolaldehyde and 3.5 mM formate production from 20 mM methanol in 1.5 h, containing 0.2 g/L PcAOX, 10 g/L GALS and 300 U/mL CAT.

**Fig. 2.**
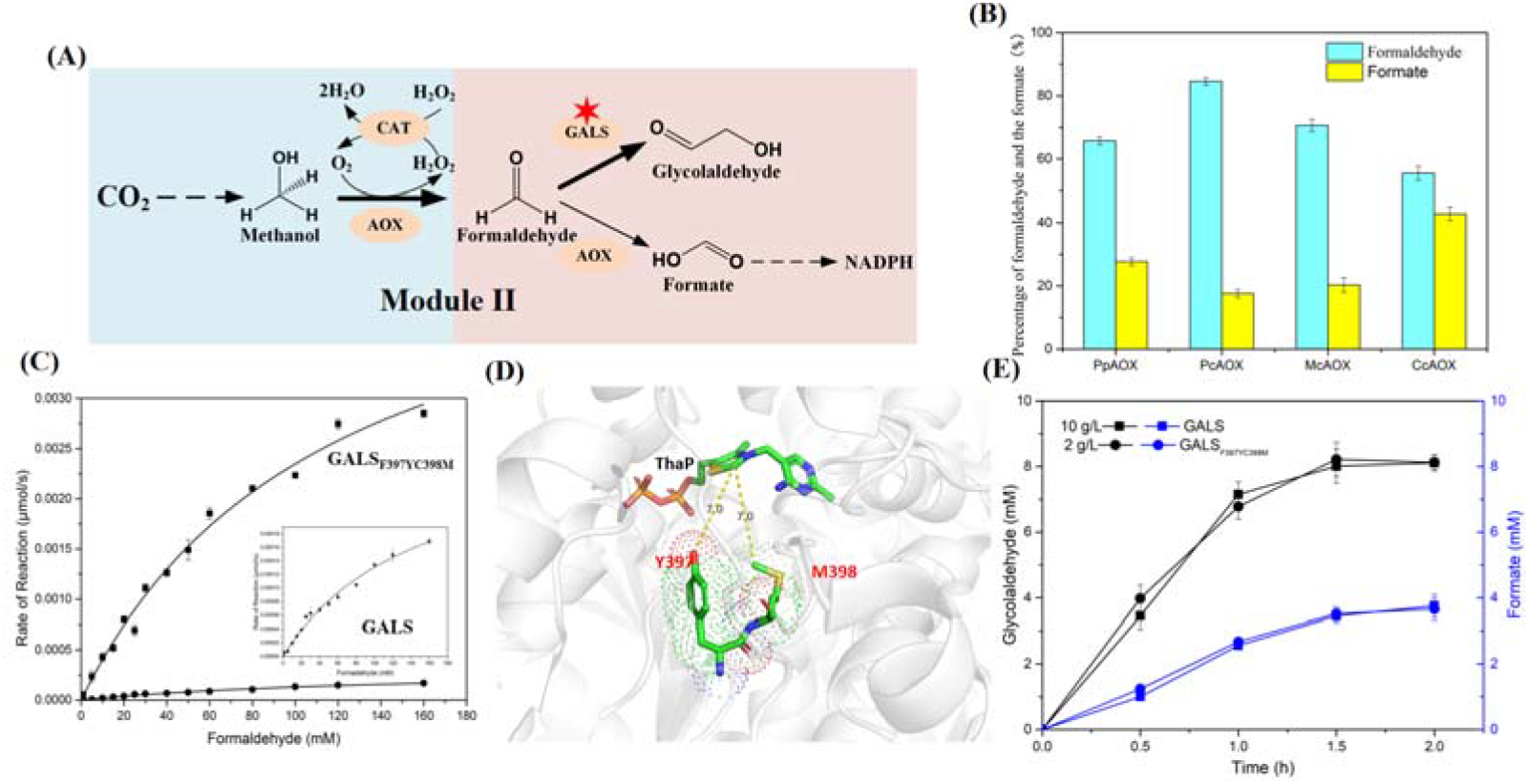
Dynamic modulation of interface from methanol to glycolaldehyde. **(A)** Schematic of module II from CO_2_ to glycolaldehyde and formate, with individual modules colored. All enzymes and chemicals are indicated. Red asterisk indicates key elements to be improved. **(B)** Screen of methanol oxidase (AOX). The reaction mixture (0.2 mL) was performed under the condition of 20 mM methanol, at 37□, 1.5 h. **(C)** Enzyme kinetics of GALS and mutant. The main figure is enzyme kinetics of GALS and mutant. Enlarged view show enzyme kinetics of GALS more accurately. Enzyme kinetics were determined with 0.05 mg mL^-1^ enzyme. The concentration of formaldehyde ranged from 0 to 160 mM. **(D)** The overview of the selected two mutations in active center. The yellow dotted lines indicate distances between the ThaP and two mutations. **(E)** Time profiles of glycolaldehyde and formate production under the condition of 10 g/L GALS or 2 g/L GALS variant. The reaction mixture (0.5 mL) was performed under the condition of 20 mM methanol, at 37 □, 1.5 h. All values shown are means of triplicate measurements. The error bars represent standard deviations.

The GALS is a key element in the hybrid system, however it accounted for 90% of the total protein dosage in the module II. In order to cut down the concentration of total protein, we determined to improve the catalytic activity of GALS. We proposed to screen significant active site residues around the active center, where 14 previously identified positions were selected to do single-point saturation mutagenesis[10]. The sub-saturating concentrations of formaldehyde (30 mM) was employed to screen for GALS variants with higher affinity for substrate and improved activity for glycolaldehyde production. After screening, libraries of positions N27,E28,F397 and C398 contained more variants with significantly increased activities (Supplementary Fig. 3). Subsequently, we introduced a four-site random combination of mutations into GALS and selected the highest active mutant. After screening of more than 5000 clones the beneficial combinations F397Y and C398M were identified and the double mutant (GALS_F397YC398M_) enabled the substrate affinity and catalytic efficiency to reach 117 mM and 146 M^-1^S^-1^, respectively, where the kcat of variant was improved approximately 10.8-fold than GALS (Fig. 2c). The mutations of F397Y and C398M are all located in the monomer-monomer interface, and the substitutions shrinked the volume of substrate binding pocket and may enhanced the interaction between ThDP and substrate (Fig. 2d). Finally, 2 g/L GALS_F397YC398M_ produced the similar yield of glycolaldehyde with 10 g/L GALS (Fig. 2e). The total protein dosage was reduced by approximately 5-fold, which was superior for industrial scale-up and commercialization.

### Precise optimization of module III

Subsequently, the module III from glycolaldehyde to PHB was optimized containing six enzymes: acetyl-phosphate synthase (ACPS), phosphate acetyltransferase (PTA), Acetyl-CoA acetyltransferase (PhaA), Acetoacetyl-CoA reductase (PhaB), PHB synthase (PhaC) and formate dehydrogenase (FDH) (Fig. 3a). In order to eliminate by-product acetate, we balanced the reaction rates by fine-tuning the ratio of individual enzyme dosages to prevent the accumulation of intermediates. The increases in all enzyme loadings by 2-fold did not significantly enhance PHB molar yield of carbon, indicating that these enzyme loadings was sufficient (Supplementary Fig. 4). To avoid the accumulation of acetyl-phosphate, the dosages of ACPS and PTA were gradually regulated to balance the flux ratio of acetyl-phosphate synthesis and consumption. The enzyme dosages of the PHB synthesis pathway were also adjusted to pull fluxes into the downstream reactions. By rerouting the carbon metabolic flux, enzymatic reaction rates in the cascade reached an equilibrium with only slight amounts of acetate (Fig. 3b).

**Fig. 3.**
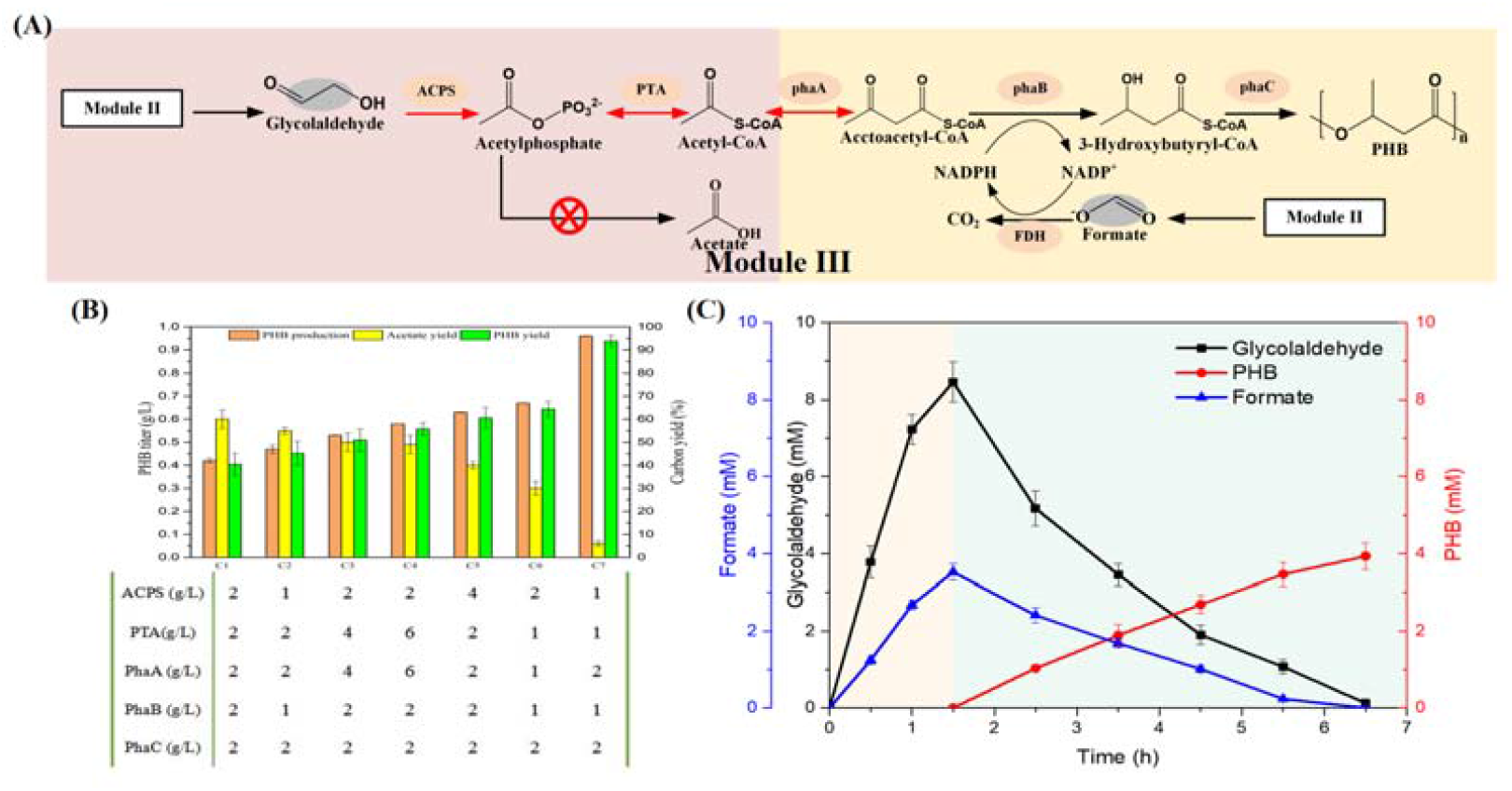
Optimization of pathway from glycolaldehyde to PHB. **(A)** Schematic of module II from CO_2_ to glycolaldehyde and formate, with individual modules colored. All enzymes and chemicals are indicated. The grey indicates products from module II. The red cross indicates deletion of by-product pathway. **(B)** Optimization of enzyme loadings for balancing the flux ratio of pathway. The forms described main enzyme loadings in this pathway. The reaction mixture (0.5 mL) was performed under the condition of 20 mM glycolaldehyde, at 37 □, 5 h. **(C)** Demonstration of the integrated pathway from methanol to PHB. The reaction mixture (0.5 mL) was performed under the condition of 20 mM methanol as initiated substrate, at 37 □, 5 h. All values shown are means of triplicate measurements. The error bars represent standard deviations.

Finally, 20 mM methanol was used as a substrate for PHB synthesis by combing module II and III. Firstly, the methanol was converted into 8.46 mM glycolaldehyde and 3.5 mM formate in 1.5 hour. After removing enzymes from this system, approximately 4 mM of PHB was accumulated in the subsequent 5 hours by supplementing the remaining six enzymes and auxiliary components (Fig. 3c). This system achieved a carbon molar yield of approximately 93.6%, exceeding all PHB biosynthetic pathways reported to date [18, 19].

### PHB synthesis via hybrid system from CO_2_

Based on the above optimization, we attempted to *de novo* synthesize PHB from CO_2_ and hydrogen by coupling the enzymatic processes with CO_2_ reduction (Fig. 4a). In fact, as increasing substrate concentration, module III was capable of maintaining high carbon molar yield up to 300 mM glycolaldehyde (Supplementary Fig. 5a). Unfortunately, the yield of glycolaldehyde began to decrease when over 20 mM methanol was added in module II(Supplementary Fig. 5b). It should be ascribed to the specificity of the AOX enzyme for glycolaldehyde and insufficient dissolved oxygen in the system. To further test the scale up potential of this system, scaling-up technology of one-pot concentration was introduced in subsequent projects.

**Fig. 4.**
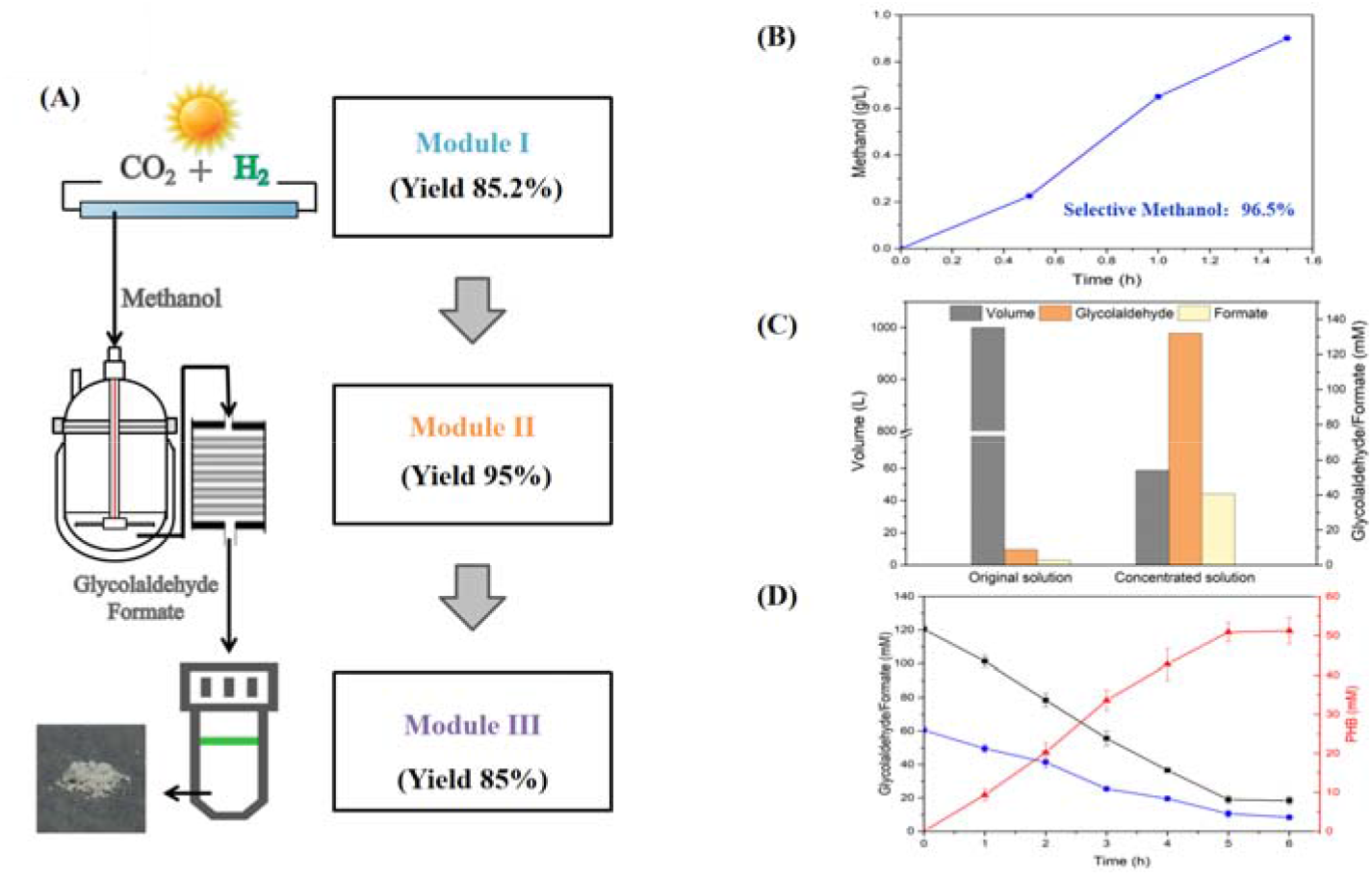
PHB synthesis via hybrid system from CO_2_. **(A)** The schematic procedure of PHB synthesis from CO_2_ and hydrogen. Yield of module I indicates molar conversion efficiency of CO_2._ Yield of module II indicates carbon yield of enzymatic reaction and loss efficiency of product in the physical enrichment process. Yield of module III indicates carbon yield of last enzymatic reaction. **(B)** Time profiles of methanol production. The reaction was operated at 250° C, 5 MPa. The airspeed was 4000 h^-1^. **(C)** Effect of physical enrichment process. The reaction solution was collected centrally and then was concentrated by Vacuum Concentrator at ambient temperature and pressure. **(D)** Time profiles of PHB production and glycolaldehyde and formate consumption in the last enzymatic process. The reaction mixture (1 mL) was performed under the condition of processed solution, at 37 □, 6 h. All values shown are means of triplicate measurements. The error bars represent standard deviations.

Firstly, the chemical reaction unit was operated at 250° C, 5 MPa and CO_2_ was chemically hydrogenated to methanol at a rate of 18 mM h^-1^ g^-1^ Cu-based catalyst with molar conversion of carbon of 85.2% and hydrogen of 84.8% (Fig.4b)[20]. The produced methanol was constantly condensed and fed into the enzymatic unit to a final concentration of 20 mM. In the enzymatic reaction unit, a 1 L volume of glycolaldehyde synthesis module was carried out, yielding approximately the equivalent glycolaldehyde titer and efficiency as the micro-scale system system in 1.5 hours. After stopping the reaction and removing enzyme preparation, the reaction solution was concentrated by vacuum enrichment at ambient temperature and pressure. Finally, the volume of 1 L is concentrated to 58.8 ml and a 5% loss of target product was detected in the physical treatment process (Fig.4c).

The processed solution was added as a substrate into subsequent module as well as supplementing the remaining enzymes and cofactors. The last enzymatic reaction produced 5.8 g/L PHB in total in the subsequent 5 hours, corresponding to a slightly dropped carbon yield (85%) (Fig.4d). Compared to other *in vitro* biosynthetic systems for PHB production, this artificial hybrid system features higher efficiencies in enzymatic reaction. The excellent molar conversion efficiency of carbon was benefited from design of atom-economic pathway and efficient engineered enzymes. By using spatial and temporal segregation of steps, the hybrid system achieved a PHB productivity of 1.06 g^-1^L^-1^h^-1^ from CO_2_ with 73.6% of molar utilization efficiency of CO_2_, exceeding that of other PHB synthesis biosystems from CO_2_.

## Discussion

Key challenges of synthesis of PHB based on CO_2_ are powerful energy efficiency and efficient carbon-conserving pathway for carbon-negative biosynthesis. The hybrid system is the most appealing strategy due to its high energy efficiency and stability[21]. Recently, chemoautotrophic photo-electrosynthesis have been gaining prominence for fixing CO_2_ and some typical electro-biological models are reported. A hybrid inorganic-biological system in combination with the *Ralstonia eutropha* drove CO_2_ direct fixation into PHB, resulting in a titer of 700 mg/l and efficiency (η_elec_) of 36%[22]. Another integrated two-step process was developed, coupling the hybrid CO_2_ electrosynthesis with acetate fermentation by *Sporomusa ovata* and *Cupriavidus basilensis*, which converted CO_2_ to PHB with 11.06% of overall carbon molar yield[23]. The solar energy conversion rate of the chemoautotrophic photo-electrosynthesis system is about 7-8%, which is a significant increase compared to the photosynthetic system[24]. However, commercial scale is still not available, as electron transfer mechanism and electrode-biological interface remain issues to be resolved. Moreover, the CO_2_ assimilation pathway in autotrophic systems were mainly dependent on the CBB cycle and Wood-Ljungdahl pathway, so the carbon yield of PHB was limited by CO_2_ emission via natural carbon metabolisms in host strain.

Over the recent years, CO_2_ hydrogenation to methanol is one of the attractive and potentially profitable routes in CCSU (Carbon Capture, Storage, and Utilization)[25]. Bio-manufacturing based on artificial atomic economic pathway using green methanol as a substrate offers carbon-negative route to synthesize target products in environmentally friendly manner. Very recently, this approach has been demonstrated for starch synthesis from directly CO_2_ via 11 core reactions.[26]. In this work, we described a bio-hybrid system, coupling energy capture with carbon fixation and conversion, that efficiently converts CO_2_ to PHB, resulting in 5.8 g/L with 73.6% of CO_2_ utilization yield and maximum 93.6% of carbon molar yield in biological process. This hybrid system displayed much higher carbon yield and conversion rate than the chemoenzymic system for starch. We believe that the integration of photovoltaic solar-harvesters, chemically reduced CO_2_ and enzymatic biocatalysts would representing a promising carbon-neutral way to fundamentally solve the CO_2_ utilization.

Plastic pollution worldwide has raised the demand for biodegradable bioplastics[27, 28]. PHAs is considered as a promising alternative to traditional chemical plastics because it rapidly and completely degrades in environment[29]. However, although PHAs have been extensively studied for 30 years, their commercialization is limited due to high cost of the production procedure[30]. The PHA produced by current microbiological fermentation has resulted in a high price of US$4-6/kg, 5-6 fold that of petroleum-based plastics[30]. Reducing the cost of raw materials is the essential key to overcome this issue. In our hybrid system, the CO_2_ consumption for obtaining 1 kg of methanol is 1.4 kg and the hydrogen consumption is 0.2 kg. The integrated costs of 1 kg of methanol is approximately US$0.37, as measured by the photovoltaic power guide price and CO_2_ capture costs in CCUS (Carbon Capture, Storage, and Utilization). The raw material cost of 1kg of PHB is approximately US$0.67, which is only 1/6-1/10 of the current price. This price is expected to be continuously reduced because photovoltaic electricity prices will continue to plummet with cost decrease in renewable technologies and local policy supports.

Current global CO_2_ emissions are already 37 billion tonnes per year and global atmospheric CO_2_ concentration is expected to reach 500 ppm by 2045[31]. The exacerbating climate crisis has accelerated the demand for carbon-negative chemical manufacturing and renewable economic models[32]. A carbon-negative manufacturing for PHB allows the establishment of a complete closed-loop PHB production from CO_2_ and its degradation to CO_2._ We envisioned a practical prospect of 3G bio-manufacturing platform that artificial hybrid system drove efficient carbon-neutral manufacturing. In the future, the key will be to keep enzyme costs low by finding stable enzymes so that they can be used for long periods of time, creating methods for recycling the enzymes, and developing inexpensive purification methods. These are largely technical rather than fundamental challenges. We therefore propose further development of hybrid system for PHB synthesis from CO_2_.

## Conclusion

In this study, we developed an artificial hybrid system to produce bio-degradable plastic PHB directly from CO_2_. A space-coupled hybrid system with excellent carbon fixation rate and molar conversion efficiency was constructed. In the hybrid process, carbon molar yield of 93.6% was achieved for producing PHB from methanol. Overall, this study provides a feasible alternative solution to address plastic pollution and excessive CO_2_ emissions simultaneously.

## Acknowledgments

We thank the core facility center at Tianjin Institution of Industrial Biotechnology, CAS, for instrument and technology support.

## Funding

National Key R&D Program of China Grant 2021YFC2103500 (YWL)

National Natural Science Foundation of China NSFC-32001030

Strategic Priority Research Program of the Chinese Academy of Sciences-Precision

Seed Design and Breeding grant XDA24020103-3 (HFJ)

Tianjin Synthetic Biotechnology Innovation Capacity Improvement Project Grant TSBICIP-KJGG-007 (HFJ)

## Author contributions

Conceptualization: HFJ, YHM, JZ, DYL

Methodology: HFJ, JZ, DYL

Investigation: JZ, DYL, YWL, HYC, JC, HDZ, SPF

Visualization: HFJ, JZ, DYL

Funding acquisition: YWL, DYL, HFJ, YHM

Project administration: HFJ

Supervision: HFJ, YHM

Writing – original draft: DYL, JZ, HFJ

## Data and materials availability

All data are available in the main text or the supplementary materials.

## Supplementary Materials

Materials and Methods

Figs. S1 to S5

Tables S1 to S4

References 1-5

