## Supplemental materials for "Hybrid Synthesis of bioplastics polyhydroxybutyrate from carbon dioxide"

**Materials and Methods**

**Chemicals and agents**

Common chemicals were bought from Sigma-Aldrich (Shanghai, China), SolarBio (Beijing, China), Zhenzhun Biotech (Shanghai, China) and Yuanye Biotech (Shanghai, China). Standard acetic acid, formaldehyde (FALD), glycoaldehyde (GALD), dihydroxyacetone (DHA), and acetyl phosphate (AcP) were purchased from Yuanye Biotech (Shanghai, China). Restriction enzymes and DNA polymerase were purchased from Thermo Fisher Scientific (Shanghai, China), and TransGen Biotech (Beijing, China). Kits for DNA manipulation were purchased from Axygen (Shanghai, China), BL21(DE3) and DH5α cells were purchased from TransGen Biotech (Beijing, China). Primers and synthesized genes were obtained from Genecreat (Wuhan, China) or GENEWIZ (Suzhou, China). Materials and equipment for protein purification were obtained from GE Healthcare (Beijing, China) and BioRad (Beijing, China). HPX-87H column were purchased from BioRad (Beijing, China).

**Plasmids construction**

Plasmids used in the study are listed in Supplementary Table 4. The plasmids of mutagenesis libraries were constructed by Gibson DNA assembly. The plasmids for protein expression were constructed into the pET28a plasmid.The genes were inserted between the *NdeІ* and *XhoI* restriction sites.There was an 6×His tag in N-terminal for purification.

**Bacterial strains and growth condition**

*E. coli* BL21 (DE3) (TransGenTM) was grown at 37 ºC or 16 ºC in 2YT medium for protein expression. Antibiotics for selection purposes were used accordingly at 50 μg ml^-1^ kanamycin.

**Purification of alcohol oxidases**

Transformed *E. coli* BL21(DE3) was grown in 2YT medium at 37 °C. Protein expression was induced when the cultures reached OD_600_≈0.6-0.8 by adding 0.5 mM isopropyl β-D-1-thiogalactopyrano(IPTG). Next, the cells were incubated at 16 °C until late stationary phase and then harvested by centrifugation at 5500 rpm for 10 min at 4 °C. Cells were resuspended in 40 ml lysis buffer (50 mM potassium phosphate pH 7.8, 400 mM NaCl, 100 μM FAD).The high-pressure homogenizer cracked the bacteria to release the proteins. After the removal of cellular debris by centrifugation (10000 rpm, 4 °C, 60 min), the supernatant was loaded onto His-Spin protein mini-prep columns pre-equilibrated using the lysis buffer .The elution of the proteins was performed using a 50−400 mM imidazole gradient. Fractions containing the pure proteins as indicated by SDS-PAGE were pooled and then desalted and concentrated using 30 kDa centrifugal filter units and 50 mM potassium phosphate buffer (pH 7.5).

**Other protein synthesis and purification**.

Catalase was purchased from Aladdin. All genes were transformed into *E. coli* BL21 (DE3) for expression and cultured with 800 ml 2YT medium. 0.5 mM IPTG was added and induced for 16h. Bacteria was collected at 5500 rpm and resuspend in pre-cooled buffer. The high-pressure homogenizer cracked the bacteria to release the proteins. Proteins were purified by His-Spin protein mini-prep columns (Zymo Research). 50 mM and 100 mM imidazole was used to wash out impurities.Then used 200mM imidazole to elute target proteins. Use Amicon Ultra-15 ultrafiltration tube to concentrate proteins. The proteins concentration was determined using a BCA Protein Assay Reagent Kit (Pierce, USA) with 2 mg/ml BSA as the standard.

**Demonstration of the module III in vitro.**

Glycolaldehyde as the initial substrate: The assay was set up in a final volume of 0.5 mL at 37 °C, 5 h, containing 50 mM HEPES buffer (pH 7.5), 5 mM MgSO_4_, 5 mM K_3_PO_4_,1 mM ThDP, 0.5 mM CoA, 0.5 mM NADP^+^, 2 mg·mL^−1^ ACPS, 1 mg·mL^−1^ PTA, 2 mg·mL^−1^ PhaA,1 mg·mL^−1^ PhaB, 2 mg·mL^−1^ PhaC, 2 mg·mL^−1^ FDPH. The different enzyme loadings was shown as in Fig. 3b.

**The complete process in vitro from methanol to PHB.**

Firstly, methanol as the initial substrate, the assay was set up at 37 °C in a final volume of 0.5 mL containing 50 mM HEPES buffer (pH 7.5), 5 mM MgSO_4_, 1 mM ThDP, 0.2 mg·mL^-1^ AOX, 300U mL^-1^ CAT, 2 mg·mL^-1^ GALS_F397YC398M_ (or 10 mg·mL^-1^ GALS). After 1.5 h, The reaction solution was fed into micro-ultrafiltration tube (3 KDa) to intercept proteins, at 3500 rpm, 0.5 h. The filtrate was used as the initial substrate for next step.

The second stage was initiated by supplementing the remaining enzymes and auxiliary components, containing 5 mM K_3_PO_4_, 0.5 mM CoA, 0.5 mM NADP^+^, 2 mg·mL^−1^ ACPS, 1 mg·mL^−1^ PTA, 2 mg·mL^−1^ PhaA,1 mg·mL^−1^ PhaB, 2 mg·mL^−1^ PhaC, 2 mg·mL^−1^ FDPH. The reaction volume is filled to 0.5 mL by HEPES buffer (pH 7.5), 5 mM MgSO_4_, 5 mM K_3_PO_4_. The assay was set up at 37 °C for 5 h.

**Assay of formaldehyde:**

Preparation of 0.25% acetylacetone solution: weigh 25 g of ammonium acetate and soluble in a little water, then add 3 ml glacial acetic acid and 0.25 ml acetylacetone, after mixing, make up to 100 ml and adjust the pH to 6.0.

Detection of formaldehyde: take 50 ul of sample and add 150 ul acetylacetone solution to reaction at 60 ℃ for 10 min，then detect the absorbance value at 414 nm. The amount of formaldehyde was calculated from the standard curve.

**Assay of formate,glycoaldehyde, acetate and acetyl-phosphate.**

HPLC detection for formate: formate was detected by HPLC. HPLC conditions: column, Aminex HPX-87H (Bio-Rad); detection wavelength, 210 nm; mobile phase, 5 mM sulphuric acid; flow rate, 0.6 mL·min-1; sample volume, 20 µL; column temperature, 40 ℃.

HPLC detection for glycoaldehyde: glycoaldehyde was detected by HPLC. HPLC conditions: column, Aminex HPX-87H (Bio-Rad); detection wavelength, 210 nm; mobile phase, 5 mM sulphuric acid; flow rate, 0.6 mL·min^-1^; sample volume, 20 µL; column temperature, 40 ℃.

HPLC detection for acetate: acetate was detected by HPLC. HPLC conditions: column, Aminex HPX-87H (Bio-Rad); detection wavelength, 210 nm; mobile phase, 5 mM sulphuric acid; flow rate, 0.6 mL min-1; sample volume, 20 µL; column temperature, 40 ℃

HPLC detection for acetyl-phosphate: add equal volume of 5% sulfuric acid to the sample for completely decomposing AcP into acetic acid. Acetic acid was then detected by HPLC. HPLC conditions: column, Aminex HPX-87H (Bio-Rad); detection wavelength, 210 nm; mobile phase, 5 mM sulphuric acid; flow rate, 0.6 mL min-1; sample volume, 20 µL; column temperature, 40 ℃.

**Analytical method of PHB**

Samples after reaction were centrifuged and dried before digestion in 99.99% sulfuric acid for 30 min at 95 ℃. The acid-digested samples were then allowed to cool to room temperature for 30 min prior to filtering the samples through a 0.2 µm PVDF syringe filter. The content of PHB was detected by high performance liquid chromatography (HPLC) equipped with an HPX-87H column.HPLC conditions: detection wavelength, 210 nm; mobile phase, 5 mM sulphuric acid; flow rate, 0.6 mL·min^-1^; sample volume, 20 µL; column temperature, 40 ℃. Different weights of PHB standards were processed in the same way and calculated to obtain a standard curve.

**Screening candidates of Alcohol oxidases**

The recently described alcohol oxidase from the white-rot basidiomycete Phanerochaete chrysosporium (PcAOX) was reported to express in *E coli*[4]. We screened similar species from related evolutionary tree and selected 10 genes to synthesis in GenScript(Nanjing,China) .

**Iterative Saturation Mutagenesis of GALS**

The determination of glycolaldehyde is as follows: 30 μL different concentrations of glycolaldehyde were prepared, and then 150 μL spetrophotometric chromogenic reagent (1.5 g diphenylamine was dissolved into 100 mL acetic acid, then added 1.5 mL concentrated sulfuric acid) was added, keeping at 90 °C for 15 min. At last, product concentration was measured by spectrophotometrically monitoring at 652 nm.

In order to obtain the desired saturation mutagenesis, oligonucleotide primers were designed with degenerate codon NNK. For 95% library coverage, the screening of 96 transformants for single-site saturation mutant was required by using NNK codon degeneracy. Each single-site saturation mutant library was generated according to the PCR-based Quick Change method. PCR reaction was performed with Fastpfu DNA Polymerase (Transgen, China) under the following conditions: the reaction was started at 94 °C (5 min), followed by 30 cycles 94 °C (20 s), 58 °C (20 s), 72 °C (3.5 min), with a ﬁnal extension at 72 °C (5 min). The PCR product was digested with *Dpn*I restriction enzyme and transformed into *E. coli* BL21 (DE3) competent cells to create the library for screening.

Each of the mutant colonies was picked and incubated 24 hours in 200 μL LB medium with 100 μg mL^-1^ kanamycin while shaking at 37 °C in 96-well microplate, and then scaled up to 1 mL LB medium for [protein expression](D:/Users/SunTao/AppData/Local/Youdao/Dict/7.5.2.0/resultui/dict/javascript:;) as well as BFD. The cell pellets were harvested by centrifugation at 3,300 g for 1 min and lysed by the re-suspension in 150 μL lysis buffer with 1 U DNase I and 1 mg mL^-1^ lysozyme, followed by 1 hour at 37 °C. Subsequently, 150 μL lysis buffer with 30 mM L^-1^ formaldehyde was added directly to the crude lysates for condensation assay and the plates were further incubated at 37 °C at 750 rpm for 90 min. After removing cells by centrifugation, 30 μL of the samples was used for a coloration assay.

**Kinetic properties of glycolaldehyde synthase**

An initial continuous assay included 50 mM potassium phosphate buffer (pH 7.4), 5 mM MgSO_4_, 0.5 mM ThDP, 50 μg mL^-1^ glycerol dehydrogenase, 1 mM NADH, and different concentrations formaldehyde. The reaction was initiated by the addition of purified BFD or mutants at 37 °C, and then an initial linear decrease in absorbance at 340 nm was observed. Enzyme kinetics were determined with formaldehyde as substrate. The concentrations ranged from 0 to 160 mM. Kinetic parameters *k_cat_* and *K_m_* were estimated by measuring the initial velocities of enzymic reaction and curve-ﬁtting according to the Michaelis-Menten equation, using GraphPad Prism 5 software. All experiments were conducted in triplicate.

**Four-site combination of mutations**

Two double-sites saturation mutant library (N27, E28 and F397, C398)was constructed by consecutive degenerate codon NNK. Two rounds of directed evolution were implemented. In each round, the best double-sites mutations was obtained. Then, four beneficial substitutions were randomly fixed by single-point mutation. Finally, a library of 16 mutants was screened for the best mutants.

**The computational analysis of glycolaldehyde synthase**

To analyze the structure changes for each mutation of GALS, the backrub module in Rosetta[1] suite was used to predict the structures of the best mutant with cofactor ThDP of each round. Then the POVME[2] package was used to calculate the volume changes of the binding pocket. The backrub option file can be found in supplementary parameters.options files. And POVME configuration file can be found in POVME_protocol.ini.

**Concentration of reaction solution**

The reaction solution was fed into parallel to several ultrafiltration tubes to intercept proteins. The filtrate was collected centrally and then was concentrated by Vacuum Concentrator at ambient temperature and pressure. Precipitated salts were removed during the concentration process.

**Assay oxidative capacity of AOX and CAT for NADPH**

The oxidative ability to NADPH is demonstrated by the reduction value of OD_340._ The reaction was carried out in a 200 μL system contains 50 mM potassium phosphate buffer (pH 7.5),1 U/mL AOX or CAT, 2 mM NADPH.

**Supplementary Figure 1**

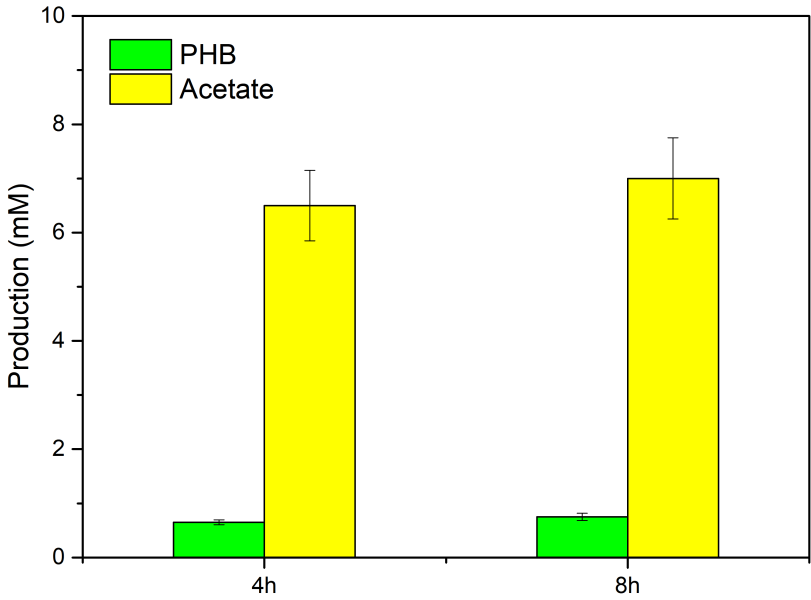

**Supplementary Figure 1. One-pot enzymatic synthesis of PHB in vitro.** The reaction mixture (0.5 mL) was performed under the condition of 20 mM methanol, at 37℃, 4 h and 8 h. The enzyme loading was 2 g/L each and the glucose-6-phosphate was supplemented to regenerate reduced nicotinamide adenine dinucleotide phosphate (NADPH) by glucose-6-phosphate-dehydrogenase (G6PD).All values shown are means of triplicate measurements. The error bars represent standard deviations.

**Supplementary Figure 2**

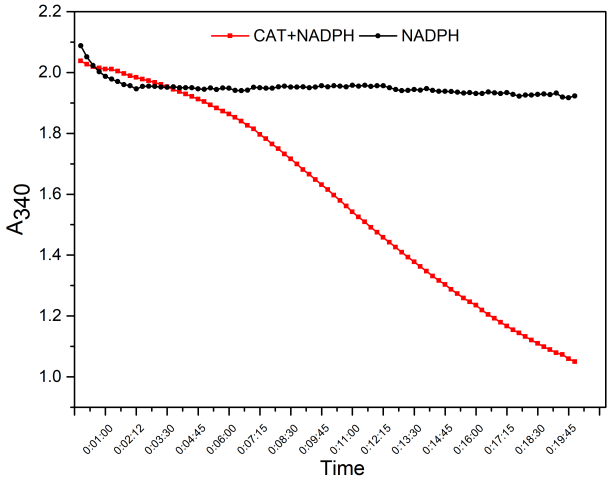

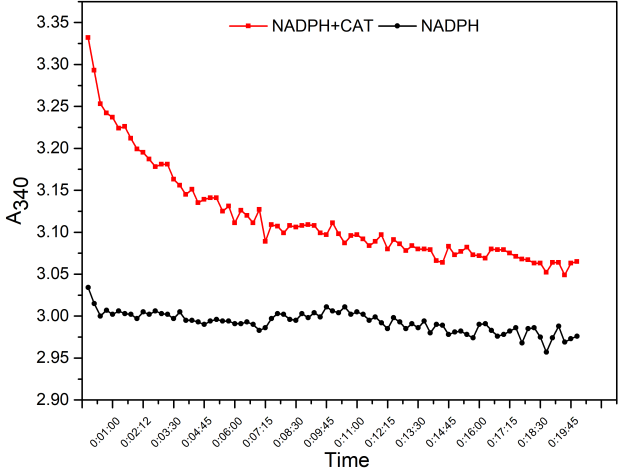

**Supplementary Figure 2. Identified the activity of AOX and CAT in response to NADPH.** The oxidative ability to NADPH is demonstrated by the reduction value of OD_340._ The reaction was carried out in a 200 μL system contains 50 mM potassium phosphate buffer (pH 7.5),1 U/mL AOX or CAT, 2 mM NADPH.

**Supplementary Figure 3**

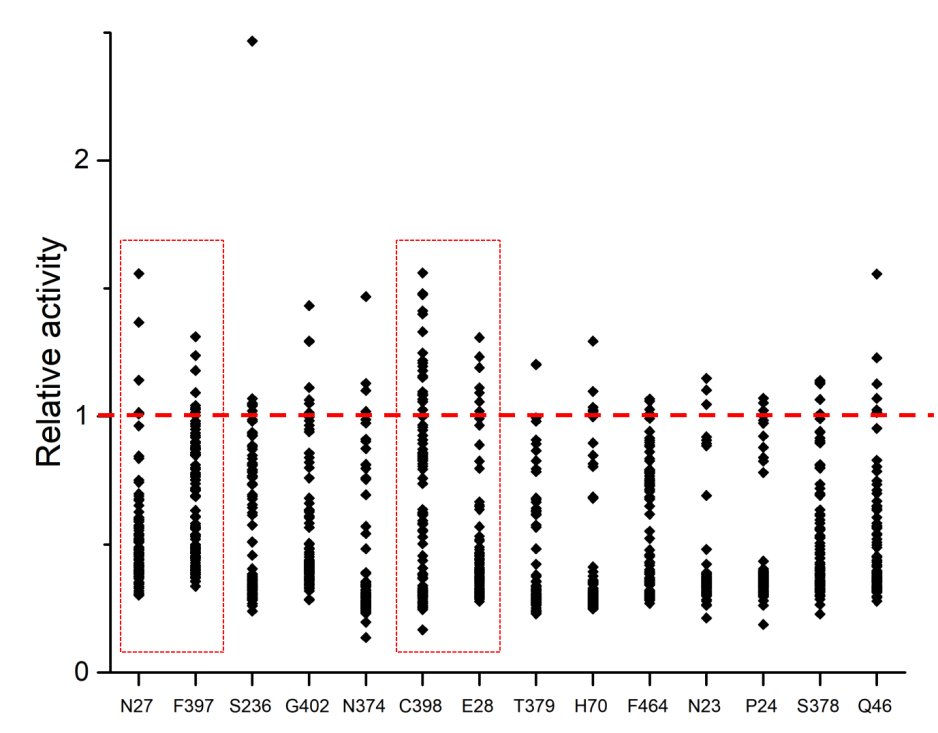

**Supplementary Figure 3. Experimental validation of single-point saturation mutant**. The x-axis label represents the selected positions in GALS. The single-point saturation mutation assays were carried out for each selected position. The y-axis label represents the relative catalytic activity of different mutants. The relative activity was defined as the ratio of the production of glycolaldehyde in the mutants to that in GALS. The yields of glycolaldehyde were determined by the chromogenic reaction. 4 positions containing higher activity mutants were for selected in the red rectangle.

**Supplementary Figure 4**

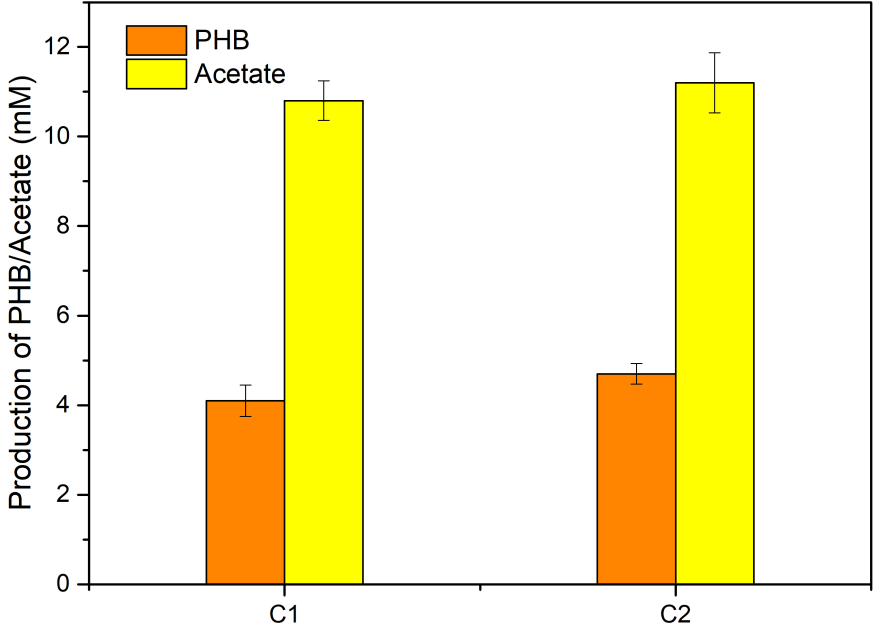

**Supplementary Figure 4. PHB production from glycolaldehyde by increasing enzyme loadings.** The reaction mixture (0.5 mL) was performed under the condition of 20 mM glycolaldehyde, at 37℃, 5 h. Group C1 indicates 2 g/L of enzyme loading in this pathway. Group C2 indicates 4 g/L of enzyme loading in this pathway. All values shown are means of triplicate measurements. The error bars represent standard deviations.

**Supplementary Figure 5**

（A） **（B）**

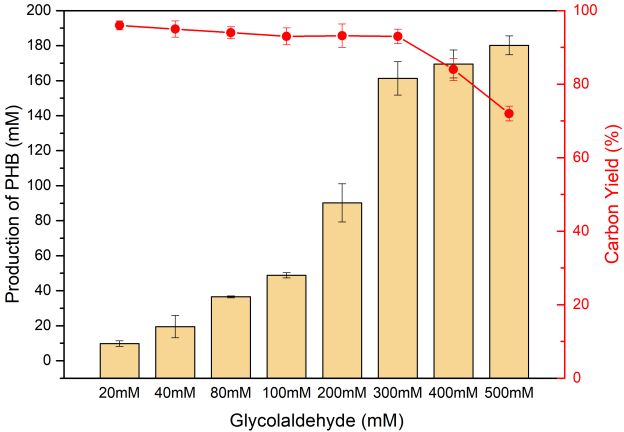

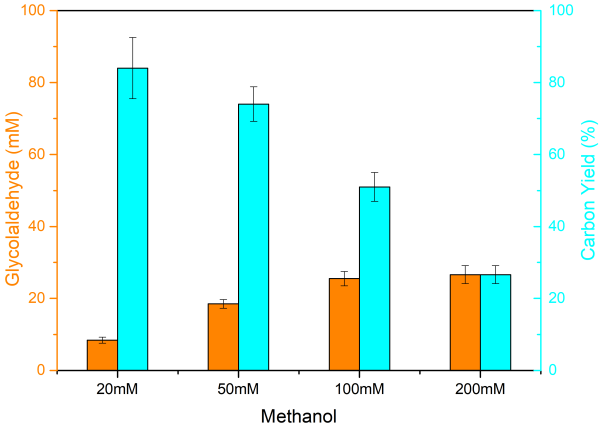

**Supplementary Figure 5. Increasing substrate concentration for module II and module III.（A）**Increasing glycolaldehyde concentration for module III. The reaction mixture (0.5 mL) was performed under the condition of 20-500 mM glycolaldehyde, at 37℃, 5 h. **(B)** Increasing methanol concentration for module II. The reaction mixture (0.5 mL) was performed under the condition of 20-200 mM methanol, at 37℃, 1.5 h. All values shown are means of triplicate measurements. The error bars represent standard deviations.

**Supplementary Table 1. Enzymes used in this hybrid system**

| Enzyme | Name | Plasmid | Tag | Organism | references |
| --- | --- | --- | --- | --- | --- |
| AOX | Alcohol oxidases | pet28a | N‐His | *Phanerochaete chrysosporium* | [3] |
| CAT | catalase | purchased from Aladdin | | | |
| FDH | formate dehydrogenase | pet28a | N‐His | *Mycolicibacterium vaccae* |  |
| GALS | glycolaldehyde synthase | pet28a | N‐His |  | [4] |
| ACPS | Phosphoketolases | pet28a | N‐His | *Bifidobacterium* | [4] |
| PTA | phosphate acetyltransferase | pet28a | N‐His | *E. coli* |  |
| PhaA | Acetyl‐CoA acetyltransferase | pet28a | N‐His | *R. eutropha* | [5] |
| PhaB | 3‐hydroxybutyl‐CoA reductase | pet28a | N‐His | *R. eutropha* | [5] |
| PhaC | Polyhydroxybutyrate synthase | pet28a | N‐His | *R. eutropha* | [5] |

**Supplementary Table 2. The thermodynamic data of all reactions**

| **Enzyme** | **Reaction** | **△G（kJ/mol）** |
| --- | --- | --- |
| AOX | Methanol+O_2_ <=> formaldehyde+H_2_O_2_ | -98.9 ± 10.3 |
| CAT | 2H_2_O_2_ <=> 2H_2_O+O_2_ | -174.8 ± 13.2 |
| GALS | 2Formaldehyde <=> glycolaldehyde | -28.7 ± 6.4 |
| ACPS | Glycolaldehyde+phosphate <=> acetyl-phosphate | -58.8 ± 3.8 |
| PTA | Acetyl-phosphate+CoA <=> acetyl‐CoA | -8.7 ± 1.2 |
| PhaA | 2Acetyl‐CoA+CoA <=> Acetoacetyl‐CoA | 25.0 ± 1.7 |
| PhaB | Acetoacetyl‐CoA+NADPH <=> 3‐hydroxybutyl‐CoA+NADP^+^ | -13.8 ± 2.3 |
| PhaC | 3‐hydroxybutyl‐CoA <=> polyhydroxybutyrate+CoA |  |
| FDH | Formate+NADP^+^ <=> CO_2_+NADPH | -14.4 ± 6.4 |

**Supplementary Table 3. Plasmids used in this study.**

| **Plasmids** | **Relevant characteristics** | **Source** |
| --- | --- | --- |
| pET28a | pBR322 ori with *pT7*; *KanR* | Novagen |
| pET28a-*PcAOX* | pET28a vector, *Nde*І*-PcAOX-Xho*I | This study |
| pET28a-*GALS* | pET28a vector, *Nde*І*-GALS-Xho*I | This study |
| pET28a-*ACPS* | pET28a vector, *Nde*І*-ACPS-Xho*I | This study |
| pET28a-*pta* | pET28a vector, *Nde*І*-pta-Xho*I | This study |
| pET28a-*phaA* | pET28a vector, *Nde*І*-phaA-Xho*I | This study |
| pET28a-*phaB* | pET28a vector, *Nde*І*-phaB-Xho*I | This study |
| pET28a-*phaC* | pET28a vector, *Nde*І*-phaC-Xho*I | This study |
| pET28a-*FDH* | pET28a vector, *Nde*І*-FDH-Xho*I | This study |

**Supplementary Table 4. The DNA sequences of used in this study**

| **Gene** | **Sequence** |
| --- | --- |
| ***Pcaox*** | ATGGGTCACCCCGAGGAGGTCGATGTCATCGTATGCGGTGGTGGTCCTGCTGGATGCGTTGTGGCGGGCAGGCTCGCTTACGCGGACCCTACACTGAAGGTCATGCTCATTGAAGGTGGTGCCAACAACCGCGATGACCCATGGGTGTACCGCCCAGGCATCTACGTCCGCAACATGCAGAGGAACGGCATCAACGACAAGGCGACGTTCTACACCGACACCATGGCTTCTTCGTATCTCCGTGGCCGTCGCAGCATCGTCCCCTGCGCCAACATCCTCGGTGGTGGTTCTTCGATCAACTTCCAGATGTACTTTCGCGCGTCTGCTTCCGATTGGGACGACTTCAAGACCGAAGGGTGGACCTGCAAGGATCTTCTGCCTCTTATGAAGCGGCTCGAGAACTACCAGAAGCCATGCAACAACGACACCCACGGCTACGATGGTCCGATTGCCATCTCCAACGGTGGACAGATCATGCCTGTCGCCCAAGACTTCCTGAGGGCTGCCCATGCGATCGGTGTGCCATACAGCGATGATATCCAGGATCTTACCACTGCACATGGTGCTGAGATTTGGGCCAAGTATATCAACCGTCACACCGGCCGTCGTAGCGATGCTGCGACCGCATACGTGCACTCCGTCATGGACGTCCAGGATAACCTCTTCCTACGCTGCAACGCCCGCGTCAGCCGCGTTCTATTCGACGACAACAACAAGGCAGTCGGCGTAGCCTATGTCCCGTCCCGCAACCGGACACACGGCGGTAAGCTCCATGAGACCATTGTCAAGGCTCGCAAGATGGTCGTCCTCAGCTCCGGCACTCTTGGTACTCCTCAGATCCTCGAGCGCTCCGGTGTCGGCAACGGCGAGCTCCTCCGCCAACTTGGCATCAAGATCGTCAGCGACCTCCCGGGTGTTGGTGAGCAGTACCAGGACCACTACGCAACGCTGTCCATATACCGTGTCTCCAACGAGTCCATTACCACCGATGACTTCCTCCGTGGTGTCAAGGACGTGCAGCGCGAGCTTTTCACAGAGTGGGAGGTTTCGCCCGAGAAGGCTCGTCTGTCGTCCAACGCCATCGACGCTGGCTTCAAGATCCGCCCAACGGAGGAAGAGCTGAAGGAGATGGGCCCTGAGTTCAACGAACTCTGGAACCGCTACTTCAAGGACAAGCCCGACAAGCCCGTCATGTTCGGCTCCATCGTCGCTGGTGCCTACGCCGACCACACACTCCTGCCGCCCGGCAAGTACATTACGATGTTCCAATTTCTCGAGTACCCGGCGTCGCGTGGCAAGATCCACATCAAGTCGCAGAACCCCTACGTCGAGCCATTCTTCGACTCCGGCTTCATGAACAACAAGGCCGACTTTGCGCCCATCCGCTGGAGCTACAAGAAGACCCGTGAGGTCGCGCGCCGCATGGATGCATTCCGTGGTGAACTGACGTCGCACCACCCGCGCTTCCACCCTGCCTCCCCCGCGGCATGCAAGGACATTGACATCGAGACTGCCAAGCAGATCTACCCCGACGGCCTCACGGTCGGCATCCACATGGGCTCGTGGCACCAGCCGTCCGAGCCGTACAAGCACGACAAGGTCATCGAGGACATCCCCTACACCGAGGAGGATGACAAGGCTATCGACGACTGGGTCGCCGATCACGTTGAGACCACCTGGCACTCGCTCGGTACCTGCGCCATGAAGCCGCGCGAGCAGGGCGGTGTTGTCGACAAGCGCCTCAACGTCTACGGCACGCAGAACCTCAAGTGTGTTGACCTGTCGATCTGCCCCGACAACCTCGGCACGAACACCTACTCGTCTGCGCTCCTCGTCGGCGAGAAGGGCGCTGATCTCATCGCTGAGGAACTCGGCCTCAAGATCAAGACCCCGCATGCTCCCGTCCCGCACGCACCCGTCCCGACCGGCAGGCCCGCTACCCAGCAGGTCCGG |
| ***gals*** | ATGGCTTCTGTTCACGGTACCACCTACGAACTGCTGCGTCGTCAGGGTATCGACACCGTTTTCGGTAACCCGGGTTCTAACGAACTGCCGTTCCTGAAAGACTTCCCGGAAGACTTCCGTTACATCCTGGCTCTGCAGGAAGCTTGCGTTGTTGGTATCGCTGACGGTTACGCTCAGGCTTCTCGTAAACCGGCTTTCATCAACCTGCACTCTGCTGCTGGTACCGGTAACGCTATGGGTGCTCTGTCTAACGCTCGTACCTCTCACTCTCCGCTGATCGTTACCGCTGGTCAGCAGACCCGTGCTATGATCGGTGTTGAAGCTGGTGAAACCAACGTTGACGCTGCTAACCTGCCGCGTCCGCTGGTTAAATGGTCTTACGAACCGGCTTCTGCTGCTGAAGTTCCGCACGCTATGTCTCGTGCTATCCACATGGCTTCTATGGCTCCGCAGGGTCCGGTTTACCTGTCTGTTCCGTACGACGACTGGGACAAAGACGCTGACCCGCAGTCTCACCACCTGTTCGACCGTCACGTTTCTTCTTCTGTTCGTCTGAACGACCAGGACCTGGACATCCTGGTTAAAGCTCTGAACTCTGCTTCTAACCCGGCTATCGTTCTGGGTCCGGACGTTGACGCTGCTAACGCTAACGCTGACTGCGTTATGCTGGCTGAACGTCTGAAAGCTCCGGTTTGGGTTGCTCCGTCTGCTCCGCGTTGCCCGTTCCCGACCCGTCACCCGTGCTTCCGTGGTCTGATGCCGGCTGGTATCGCTGCTATCTCTCAGCTGCTGGAAGGTCACGACGTTGTTCTGGTTATCGGTGCTCCGGTTTTCCGTTACGTTTTTTACGACCCGGGTCAGTACCTGAAACCGGGTACCCGTCTGATCTCTGTTACCTGCGACCCGCTGGAAGCTGCTCGTGCTCCGATGGGTGACGCTATCGTTGCTGACATCGGTGCTATGGCTTCTGCTCTGGCTAACCTGGTTGAAGAATCTTCTCGTCAGCTGCCGACCGCTGCTCCGGAACCGGCTAAAGTTGACCAGGACGCTGGTCGTCTGCACCCGGAAACCGTTTTCGACACCCTGAACGACATGGCTCCGGAAAACGCTATCTACCTGAACGAATCTACCTCTACCACCGCTCAGATGTGGCAGCGTCTGAACATGCGTAACCCGGGTTCTTACTACTTCTGCGCTGCTGGTGGTCTGGGTTTCGCTCTGCCGGCTGCTATCGGTGTTCAGCTGGCTGAACCGGAACGTCAGGTTATCGCTGTTATCGGTGACGGTTCTGCTAACTACTCTATCTCTGCTCTGTGGACCGCTGCTCAGTACAACATCCCGACCATCTTCGTTATCATGAACAACGGTACCTACGGTATGCTGCGTTGGTTCGCTGGTGTTCTGGAAGCTGAAAACGTTCCGGGTCTGGACGTTCCGGGTATCGACTTCCGTGCTCTGGCTAAAGGTTACGGTGTTCAGGCTCTGAAAGCTGACAACCTGGAACAGCTGAAAGGTTCTCTGCAGGAAGCTCTGTCTGCTAAAGGTCCGGTTCTGATCGAAGTTTCTACCGTTTCTCCGGTTAAA |
| ***acps*** | ATGACGAGTCCTGTTATTGGCACCCCTTGGAAGAAGCTGAACGCTCCGGTTTCCGAGGAAGCTATCGAAGGCGTGGATAAGTACTGGCGCGCAGCCAACTACCTCTCCATCGGCCAGATCTATCTGCGTAGCAACCCGCTGATGAAGGAGCCTTTCACCCGCGAAGACGTCAAGCACCGTCTGGTCGGTCACTGGGGCACCACCCCGGGCCTGAACTTCCTCATCGGCCACATCAACCGTCTCATTGCTGATCACCAGCAGAACACTGTGATCATCATGGGCCCGGGCCACGGCGGCCCGGCTGGTACCGCTCAGTCCTACCTGGACGGCACCTACACCGAGTACTTCCCGAACATCACCAAGGATGAGGCTGGCCTGCAGAAGTTCTTCCGCCAGTTCTCCTACCCGGGTGGCATCCCGTCCCACTACGCTCCGGAGACCCCGGGCTCCATCCACGAAGGCGGCGAGCTGGGTTACGCCCTGTCCCACGCCTACGGCGCTGTGATGAACAACCCGAGCCTGTTCGTCCCGGCCATCGTCGGCGACGGCGAAGCTGAGACCGGCCCGCTGGCCACCGGCTGGCAGTCCAACAAGCTCATCAACCCGCGCACCGACGGTATCGTGCTGCCGATCCTGCACCTCAATGGCTACAAGATCGCCAACCCGACCATCCTGTCCCGCATCTCCGACGAAGAGCTCCACGAGTTCTTCCACGGCATGGGCTATGAGCCGTACGAGTTCGTCGCTGGCTTCGACAACGAGGATCACCTGTCGATCCACCGTCGTTTCGCCGAGCTGTTCGAGACCGTCTTCGACGAGATCTGCGACATCAAGGCCGCCGCTCAGACCGACGACATGACTCGTCCGTTCTACCCGATGATCATCTTCCGTACCCCGAAGGGCTGGACCTGCCCGAAGTTCATCGACGGCAAGAAGACCGAGGGCTCCTGGCGTTCCCACCAGGTGCCGCTGGCTTCCGCCCGCGATACCGAGGCCCACTTCGAGGTCCTCAAGAACTGGCTCGAGTCCTACAAGCCGGAAGAGCTGTTCGACGAGAACGGCGCCGTGAAGCCGGAAGTCACCGCCTTCATGCCGACCGGCGAACTGCGCATCGGTGAGAACCCGAACGCCAACGGTGGCCGCATCCGCGAAGAGCTGAAGCTGCCGAAGCTGGAAGACTACGAGGTCAAGGAAGTCGCCGAGTACGGCCACGGCTGGGGCCAGCTCGAGGCCACCCGTCGTCTGGGCGTCTACACCCGCGACATCATCAAGAACAACCCGGACTCCTTCCGTATCTTCGGACCGGATGAGACCGCTTCCAACCGTCTGCAGGCCGCTTACGACGTCACCAACAAGCAGTGGGACGCCGGCTACCTGTCCGCTCAGGTCGACGAGCACATGGCTGTCACCGGCCAGGTCACCGAGCAGCTTTCCGAGCACCAGATGGAAGGCTTCCTCGAGGGCTACCTGCTGACCGGCCGTCACGGCATCTGGAGCTCCTATGAGTCCTTCGTGCACGTGATCGACTCCATGCTGAACCAGCACGCCAAGTGGCTCGAGGCTACCGTCCGCGAGATTCCGTGGCGCAAGCCGATCTCCTCCATGAACCTGCTCGTCTCCTCCCACGTGTGGCGTCAGGATCACAACGGCTTCTCCCACCAGGATCCGGGTGTCACCTCCGTCCTGCTGAACAAGTGCTTCAACAACGATCACGTGATCGGCATCTACTTCCCGGTGGATTCCAACATGCTGCTCGCTGTGGCTGAGAAGTGCTACAAGTCCACCAACAAGATCAACGCCATCATCGCCGGCAAGCAGCCGGCCGCCACCTGGCTGACCCTGGACGAAGCTCGCGCCGAGCTCGAGAAGGGTGCTGCCGAGTGGAAGTGGGCTTCCAACGTGAAGTCCAACGATGAGGCTCAGATCGTGCTCGCCGCCACCGGTGATGTTCCGACTCAGGAAATCATGGCCGCTGCCGACAAGCTGGACGCCATGGGCATCAAGTTCAAGGTCGTCAACGTGGTTGACCTGGTCAAGCTGCAGTCCGCCAAGGAGAACAACGAGGCCCTCTCCGATGAGGAGTTCGCTGAGCTGTTCACCGAGGACAAGCCGGTCCTGTTCGCTTACCACTCCTATGCCCGCGACGTGCGTGGTCTGATCTACGATCGCCCGAACCACGACAACTTCAACGTTCACGGCTACGAGGAGCAGGGCTCCACCACCACCCCGTACGACATGGTTCGCGTGAACAACATCGATCGCTACGAGCTCCAGGCTGAAGCTCTGCGCATGATCGACGCTGACAAGTACGCCGACAAGATCAACGAGCTCGAGGCCTTCCGTCAGGAAGCCTTCCAGTTCGCTGTCGACAACGGCTACGATCACCCGGATTACACCGACTGGGTCTACTCCGGTGTCAACACCAACAAGCAGGGTGCTATCTCCGCTACCGCCGCAACCGCTGGCGATAACGAGTGA |
| ***PhaA*** | atgactgacgttgtcatcgtatccgccgcccgcaccgcggtcggcaagtttggcggctcgctggccaagatcccggcaccggaactgggtgccgtggtcatcaaggccgcgctggagcgcgccggcgtcaagccggagcaggtgagcgaagtcatcatgggccaggtgctgaccgccggttcgggccagaaccccgcacgccaggccgcgatcaaggccggcctgccggcgatggtgccggccatgaccatcaacaaggtgtgcggctcgggcctgaaggccgtgatgctggccgccaacgcgatcatggcgggcgacgccgagatcgtggtggccggcggccaggaaaacatgagcgccgccccgcacgtgctgccgggctcgcgcgatggtttccgcatgggcgatgccaagctggtcgacaccatgatcgtcgacggcctgtgggacgtgtacaaccagtaccacatgggcatcaccgccgagaacgtggccaaggaatacggcatcacacgcgaggcgcaggatgagttcgccgtcggctcgcagaacaaggccgaagccgcgcagaaggccggcaagtttgacgaagagatcgtcccggtgctgatcccgcagcgcaagggcgacccggtggccttcaagaccgacgagttcgtgcgccagggcgccacgctggacagcatgtccggcctcaagcccgccttcgacaaggccggcacggtgaccgcggccaacgcctcgggcctgaacgacggcgccgccgcggtggtggtgatgtcggcggccaaggccaaggaactgggcctgaccccgctggccacgatcaagagctatgccaacgccggtgtcgatcccaaggtgatgggcatgggcccggtgccggcctccaagcgcgccctgtcgcgcgccgagtggaccccgcaagacctggacctgatggagatcaacgaggcctttgccgcgcaggcgctggcggtgcaccagcagatgggctgggacacctccaaggtcaatgtgaacggcggcgccatcgccatcggccacccgatcggcgcgtcgggctgccgtatcctggtgacgctgctgcacgagatgaagcgccgtgacgcgaagaagggcctggcctcgctgtgcatcggcggcggcatgggcgtggcgctggcagtcgagcgcaaataa |
| ***PhaB*** | atgactcagcgcattgcgtatgtgaccggcggcatgggtggtatcggaaccgccatttgccagcggctggccaaggatggctttcgtgtggtggccggttgcggccccGAAGATccgAATCAGgaaaagtggctggagcagcagaaggccctgggcttcgatttcattgcctcggaaggcaatgtggctgactgggactcgaccaagaccgcattcgacaaggtcaagtccgaggtcggcgaggttgatgtgctgatcaacaacgccggtatcacccgcgacgtggtgttccgcaagatgacccgcgccgactgggatgcggtgatcgacaccaacctgacctcgctgttcaacgtcaccaagcaggtgatcgacggcatggccgaccgtggctggggccgcatcgtcaacatctcgtcggtgaacgggcagaagggccagttcggccagaccaactactccaccgccaaggccggcctgcatggcttcaccatggcactggcgcaggaagtggcgaccaagggcgtgaccgtcaacacggtctctccgggctatatcgccaccgacatggtcaaggcgatccgccaggacgtgctcgacaagatcgtcgcgacgatcccggtcaagcgcctgggcctgccggaagagatcgcctcgatctgcgcctggttgtcgtcggaggagtccggtttctcgaccggcgccgacttctcgctcaacggcggcctgcatatgggctga |
| ***PhaC*** | atggcgaccggcaaaggcgcggcagcttccacgcaggaaggcaagtcccaaccattcaaggtcacgccggggccattcgatccagccacatggctggaatggtcccgccagtggcagggcactgaaggcaacggccacgcggccgcgtccggcattccgggcctggatgcgctggcaggcgtcaagatcgcgccggcgcagctgggtgatatccagcagcgctacatgaaggacttctcagcgctgtggcaggccatggccgagggcaaggccgaggccaccggtccgctgcacgaccggcgcttcgccggcgacgcatggcgcaccaacctcccatatcgcttcgctgccgcgttctacctgctcaatgcgcgcgccttgaccgagctggccgatgccgtcgaggccgatgccaagacccgccagcgcatccgcttcgcgatctcgcaatgggtcgatgcgatgtcgcccgccaacttccttgccaccaatcccgaggcgcagcgcctgctgatcgagtcgggcggcgaatcgctgcgtgccggcgtgcgcaacatgatggaagacctgacacgcggcaagatctcgcagaccgacgagagcgcgtttgaggtcggccgcaatgtcgcggtgaccgaaggcgccgtggtcttcgagaacgagtacttccagctgttgcagtacaagccgctgaccgacaaggtgcacgcgcgcccgctgctgatggtgccgccgtgcatcaacaagtactacatcctggacctgcagccggagagctcgctggtgcgccatgtggtggagcagggacatacggtgtttctggtgtcgtggcgcaatccggacgccagcatggccggcagcacctgggacgactacatcgagcacgcggccatccgcgccatcgaagtcgcgcgcgacatcagcggccaggacaagatcaacgtgctcggcttctgcgtgggcggcaccattgtctcgaccgcgctggcggtgctggccgcgcgcggcgagcacccggccgccagcgtcacgctgctgaccacgctgctggactttgccgacacgggcatcctcgacgtctttgtcgacgagggccatgtgcagttgcgcgaggccacgctgggcggcggcgccggcgcgccgtgcgcgctgctgcgcggccttgagctggccaataccttctcgttcttgcgcccgaacgacctggtgtggaactacgtggtcgacaactacctgaagggcaacacgccggtgccgttcgacctgctgttctggaacggcgacgccaccaacctgccggggccgtggtactgctggtacctgcgccacacctacctgcagaacgagctcaaggtaccgggcaagctgaccgtgtgcggcgtgccggtggacctggccagcatcgacgtgccgacctatatctacggctcgcgcgaagaccatatcgtgccgtggaccgcggcctatgcctcgaccgcgctgctggcgaacaagctgcgcttcgtgctgggtgcgtcgggccatatcgccggtgtgatcaacccgccggccaagaacaagcgcagccactggactaacgatgcgctgccggagtcgccgcagcaatggctggccggcgccatcgagcatcacggcagctggtggccggactggaccgcatggctggccgggcaggccggcgcgaaacgcgccgcgcccgccaactatggcaatgcgcgctatcgcgcaatcgaacccgcgcctgggcgatacgtcaaagccaaggcatga |
| ***FDH*** | CATATGGCAAAAGTGCTGTGCGTGCTGTATGATGATCCGGTGGATGGTTATCCGAAAACCTATGCACGTGATGATCTGCCGAAAATTGATCATTATCCGGGTGGCCAGATTCTGCCGACCCCGAAAGCAATTGATTTTACCCCGGGCCAGCTGCTGGGTAGTGTTAGTGGCGAACTGGGCCTGCGCGAATATCTGGAAAGCAATGGTCATACCCTGGTGGTGACCAGTGATAAAGATGGTCCGGATAGTGTGTTTGAACGTGAACTGGTGGATGCAGATGTGGTGATTAGTCAGCCGTTTTGGCCGGCCTATCTGACCCCGGAACGCATTGCCAAAGCCAAAAATCTGAAACTGGCACTGACCGCAGGCATTGGTAGTGATCATGTGGATCTGCAGAGCGCCATTGATCGTAATGTTACCGTTGCAGAAGTGACCTATAGTAATAGTATTAGCGTGGCCGAACATGTTGTTATGATGATTCTGAGCCTGGTGCGTAATTATCTGCCGAGCCATGAATGGGCACGCAAAGGTGGTTGGAATATTGCCGATTGCGTTAGTCATGCCTATGATCTGGAAGCCATGCATGTGGGCACCGTGGCCGCAGGCCGCATTGGTCTGGCCGTGCTGCGCCGTCTGGCCCCTTTTGATGTTCATCTGCATTATACCCAGCGTCATCGTCTGCCGGAAAGTGTGGAAAAAGAACTGAATCTGACCTGGCATGCAACCCGTGAAGATATGTATCCGGTGTGTGATGTGGTGACCCTGAATGTTCCGCTGCATCCGGAAACCGAACACATGATTAATGATGAAACCCTGAAACTGTTTAAGCGCGGTGCATATATTGTGAATACCGCACGTGGTAAACTGTGTGATCGCGATGCAGTGGCCCGCGCCCTGGAAAGTGGTCGTCTGGCAGGTTATGCCGGCGATGTTTGGTTTCCGCAGCCGGCACCGAAAGATCATCCGTGGCGCACCATGCCGTATAATGGCATGACCCCGCATATTAGCGGCACCACCCTGACCGCCCAGGCACGTTATGCCGCCGGCACCCGTGAAATTCTGGAATGTTTTTTTGAAGGCCGTCCGATTCGCGATGAATATCTGATTGTTCAGGGCGGTGCACTGGCCGGCACCGGTGCACATAGCTATAGTAAAGGTAATGCAACCGGTGGTAGCGAAGAAGCCGCAAAATTTAAAAAAGCCGTTCTCGAG |
